# Collateral sensitivity to β-lactam drugs in drug-resistant tuberculosis is driven by the transcriptional wiring of BlaI operon genes

**DOI:** 10.1101/227538

**Authors:** AS Trigos, BW Goudey, J Bedő, TC Conway, NG Faux, KL Wyres

**Affiliations:** IBM Research – Australia, Carlton, VIC, 3053, Australia; Sir Peter MacCallum Department of Oncology, The University of Melbourne, Parkville, VIC 3010, Australia; Cancer Research Division, Peter MacCallum Cancer Centre, Melbourne, VIC 3000, Australia; The Department of Computing and Information Systems, The University of Melbourne, Parkville, VIC 3010, Australia; Centre For Epidemiology and Biostatistics, The University of Melbourne, Parkville, VIC 3010, Australia; Bioinformatics Division, The Walter and Eliza Hall Institute of Medical Research, Parkville, Victoria, 3052, Australia; The Florey Institute of Neuroscience and Mental Health, The University of Melbourne, Parkville, VIC 3052; Department of Biochemistry and Molecular Biology, Bio21 Molecular Science and Biotechnology Institute, University of Melbourne, Parkville, Victoria 3010

**Keywords:** tuberculosis, β-lactams, antimicrobial resistance, collateral sensitivity, paradoxical hypersensitivity

## Abstract

**Background:** The evolution and spread of antimicrobial resistance is a major global public health threat. In some cases the evolution of resistance to one antimicrobial seemingly results in enhanced sensitivity to another (known as ‘collateral sensitivity’). This largely underexplored phenomenon represents a fascinating evolutionary paradigm that opens new therapeutic possibilities for patients infected with pathogens unresponsive to classical treatments. Intrinsic resistance to β-lactams in *Mycobacterium tuberculosis* (*Mtb*, the causative agent of tuberculosis) has traditionally curtailed the use of these low-cost and easy-to-administer drugs for tuberculosis treatment. Recently, β-lactam sensitivity has been reported in strains resistant to classical tuberculosis drug therapy, leading to a resurgence of interest in using β-lactams in the clinic. Unfortunately though, there remains a limited understanding of the mechanisms driving β-lactam sensitivity.

**Methods:** We used a novel combination of systems biology and computational approaches to characterize the molecular underpinnings of β-lactam sensitivity in *Mtb*. We performed differential gene expression and coexpression analyses of genes previously associated with β-lactam sensitivity and genes associated with resistance to classical tuberculosis drugs. Protein-protein interaction and gene regulatory network analyses were used to validate regulatory interactions between these genes, and random walks through the networks identified key mediators of these interactions. Further validation was obtained using functional *in silico* knockout of gene pairs.

**Results:** Our results reveal up regulation of the key regulatory inhibitor of β-lactamase production, *blaI*, following treatment with classical drugs. Co-expression and network analyses showed direct co-regulation between genes associated with β-lactam sensitivity and those associated with resistance to classical tuberculosis treatment. *blaI* and its downstream genes (*sigC* and *atpH*) were found to be key mediators of these interactions.

**Conclusions:** Our results support the hypothesis that *Mtb* β-lactam sensitivity is a collateral consequence of the evolution of resistance to classical tuberculosis drugs, mediated through changes to transcriptional regulation. These findings support continued exploration of β-lactams for the treatment of tuberculosis, particularly for patients infected with strains resistant to classical therapies that are otherwise difficult to treat. Importantly, this work also highlights the potential of systems-level and network biology approaches to improve our understanding of collateral drug sensitivity.

## Background

Collateral antimicrobial sensitivity occurs when the evolution of resistance to one or more antimicrobials directly or indirectly causes increased sensitivity to unrelated antimicrobials [1]. There are now numerous examples of this phenomenon in the literature [2, 3], and while direct mechanisms are sometimes evident based on our understanding of individual genes or pathways [4], there is a lack of knowledge to explain collateral sensitivity between drugs of unrelated function. An improved understanding of such mechanisms can inform novel treatment strategies that limit or delay the development of resistance [1].

Tuberculosis (TB) remains a significant cause of global mortality, causing an estimated 1.5 million deaths annually (WHO 2014 Global Tuberculosis Report). It can be successfully treated through combination antimicrobial therapy targeting the causal pathogen, *Mycobacterium tuberculosis* (*Mtb*). However, successful treatment is hampered by the emergence of antimicrobial resistant *Mtb*, particularly strains resistant to multiple drugs (WHO 2014 Global Tuberculosis Report).

*Mycobacterium tuberculosis* is generally considered intrinsically resistant to the β-lactams due to its production of the *BlaC* β-lactamase and the inclusion of non-classical peptidoglycan linkages in its cell wall [5, 6]. However, recent studies have reported increased β-lactam sensitivity among some clinical isolates, largely comprising multi- or extensively-drug resistant strains [7, 8], plus isolates experimentally evolved to be resistant to aminoglycosides [9]. (Multi- and extensive-drug resistance is defined by the World Health Organization as resistance to isoniazid and rifampicin, with or without resistance to other first-line drugs; and resistance to isoniazid and rifampicin, plus any fluoroquinolone, and any of the three second-line injectable drugs (amikacin, capreomycin, and kanamycin), respectively.) These findings suggest that the evolution of resistance to classical TB drugs may lead to collateral β-lactam sensitivity. The potential application of clinical regimens including β-lactams is of particular interest, due to the comparative low-cost, ease of treatment and accessibility of these drugs [10]. Although β-lactam plus β-lactamase inhibitor treatments have shown potent activity against *Mtb* in the laboratory [6], patient treatment trials have been less promising [11, 12]. Therefore, a better understanding of how resistance to the classical drugs may result in heightened β-lactam sensitivity is required to identify those patients that may benefit from β-lactam treatment.

Recent technological and algorithmic advances have facilitated the high throughput measurement of gene expression, as well as the inference and analysis of large-scale protein-protein interaction and DNA-protein interaction networks for *Mtb*, which can facilitate systems-level investigations into the transcriptional and regulatory mechanisms behind phenomena such as collateral sensitivity. However, to date there has been limited integration of network and transcriptomic analyses to understand clinically relevant system-level mechanisms in bacteria; instead such studies have focused on the identification of new genes or the associations between drugs and resistance genes (e.g. [13, 14]).

Here we describe a novel systems-level approach for the exploration of collateral β-lactam sensitivity in *Mtb*. We combine gene expression analyses with protein-protein interaction and gene regulatory network data and functional *in silico* growth simulations. Our analyses indicate that collateral β-lactam sensitivity is the result of direct transcriptional regulation between genes associated with β-lactam sensitivity and those mediating resistance to classical TB drugs. This wiring promotes the inhibition of β-lactamases as a response to drug treatment, with genes of the BlaI operon, *blaI*, *sigC* and *atpH*, playing key roles. This work is the first to demonstrate the potential of integrative computational and systems biology approaches in the understanding of the mechanisms of collateral sensitivity.

## Methods

### 1. Genes associated with β-lactam sensitivity

A list of 111 genes associated with β-lactam sensitivity (hereafter termed β-lactam^S^ genes) in *Mtb*, and two closely related species, *Mycobacterium smegmatis* and *Mycobacterium bovis*, was obtained from multiple sources [7, 15]. A full list of genes used in this study can be found in Supp. Table S1.

### 2. Genes implicated in resistance to classical TB drugs

We compiled a list of 40 genes implicated in classical TB drug resistance (hereafter termed DR genes) from The Tuberculosis Drug Resistance Mutation Database (https://tbdreamdb.ki.se/CMS/Download.aspx) (Supp. Table S2). These included genes associated with resistance to rifampicin (RIF,n = 2), isoniazid (INH, n= 22), aminoglycosides (AMI, kanamycin/captromycin/amikacin/viomycin, n= 2), streptomycin (SM,n= 3), fluoroquinolones (FLQ, n = 2), ethambutol (EMB, n = 13), ethionamide (ETH, n = 3), para-aminosalisylic acid (PAS, n = 1) and pyrazinamide (PZA, n = 1).

### 3. Expression data

*Mtb* microarray gene expression data were obtained from sample series GSE1642 [16] from the NCBI GEO database. Data were available for *Mtb* exposed to 437 treatments, including the following *in vitro* treatment conditions: classical TB drugs as single agents (isoniazid, rifampicin, amikacin, streptomycin, levofloxacin, ofloxacin, ethambutol, ethionamide, pyrazinamide) and control conditions (7H9-based growth medium without drug treatment).

We assessed the impact of classical trug treatment by comparing the variance of expression of β-lactam^s^ genes to that across all genes. Significance testing was performed by comparison to the null distribution generated by random subsampling of *Mtb* genes (n = 111 genes with 10,000 replicates), and counting the number of times we obtained a variance equal or greater than the observed value. Differential expression analysis was performed using limma [17], where differential expression was considered significant if the q-value (i.e. a p-value that has been adjusted for the False Discovery Rate (FDR) taking into account multiple testing) < 0.05 and |fold change| > 2.

To compare the strength of correlation of expression of DR genes with β-lactam^s^ genes, we exhaustively calculated Spearman ρ between the expression of each of the individual genes, generating (1) a distribution of correlation of each individual DR gene with all β-lactam^s^ genes, and (2) a distribution of DR genes with all other genes. We then used the upper quantile of the correlation magnitude (absolute value of the correlation of expression) of each of these distributions to summarise the differences in the distribution of the strength of correlation magnitude, therefore comparing the most strongly correlated DR-genes - β-lactam^S^ genes with the most strongly correlated DR -genes - non-β-lactam^S^ genes.

### 4. *Mtb* networks

We integrated molecular interaction networks from two sources: protein-protein interactions (PPIs) (22,308 interactions) from the STRING database [18], and transcription factor-target interactions experimentally obtained using chromatin immunoprecipitations [19] as a gene regulatory (GR) network (15,054 interactions). Note that although the STRING database has been traditionally considered to be solely composed of PPIs, there are a number of regulatory interactions supported by gene co-expression analysis [18]. Genes/proteins were represented as nodes and interactions were represented as edges. Only high confidence edges were analysed: PPIs with a weight greater than 700 (the cutoff suggested by STRING as being of high confidence) and statistically significant transcription factor-target gene interactions (as defined by [19]) were considered. The power-law distribution of the combined network PPI and GR was verified using igraph [20], to ensure its biological plausibility. Network visualizations were obtained using Cytoscape v3.4.0 [21].

### 5. Significance of interactions between β-lactam^S^ and DR nodes (genes/proteins) in the molecular interaction network

We assessed the significance of the interactions between gene sets (β-lactam^S^ and DR genes) using BinoX [22] a method for estimating the distribution of crosstalk expected under a random model of a given network. In this study, distribution parameters are estimated under two different permutation procedures: permutation of the node labels or edges (denoted ‘node swap’ and ‘link swap’ respectively). Once the crosstalk distribution for the given gene set(s) has been estimated, the significance of an observed degree of crosstalk can be computed. In this study, 1000 permutations were used for both methods, and significance was corrected using Benjamini-Hochberg (and hence is reported as q-values).

### 6. Random network walks to identify β-lactam^S^ nodes influenced by DR nodes

We performed random walks between all pairs of nodes in the PPI and the GR networks separately to determine the access times as an indicator for the influence of one node over another. Random walks correspond to the possible paths taken by a random walker on a network between a pair of nodes. Access times represent the ease with which information (e.g. signal transduction, gene regulation) flows from one node to another, as it is proportional to the number of connections and available paths between nodes. Simulating the random walk was unnecessary as the access time on a finite graph has an analytical solution [23] computed via eigenvalue decomposition of the edge matrices of the networks.

To assess the similarity of the access times obtained with the PPI [18] and the GR networks [19], we investigated the stability using a multivariate extension of Spearman’s ρ [24, 25]. This allows us to assess the similarity of the top-*k* access times and determine if there is a set of stable edges with low access time.

We selected pairs of nodes comprising one β-lactam^S^ node and one DR node, and narrowed down sets of pairs with small access times in either the PPI or GR network. Given the non-symmetry of access times obtained with random walks (the access time from A->B is not equal to that from B->A), we considered the results obtained in both directions independently. Cutoffs were determined from the empirical distribution: 1054.74 for the PPI in the DR genes -> β-lactam^S^ direction, 1336.58 for the β-lactam^S^ -> DR genes direction, and 1713.37 for the GR network in the DR genes -> β-lactam^S^ direction and 2741.49 in the β-lactam^S^ -> DR genes direction.

### 7. Simulating the effect on bacterial growth of double knockout β-lactam^S^ plus DR gene mutants

To identify pairs of β-lactam^S^ and DR genes whose knockout would have the largest effect on the growth of *Mtb*, we performed simulations using the iSM810 model of *Mtb* with the PROM framework [26] and the COBRA toolbox [27], which incorporate both gene-regulatory and metabolic processes to predict growth rates after double knock-down simulations. As input we used the GR network [19] and expression data described above [16].

## Results

### 1. Treatment with classical TB drugs induces the expression of β-lactamase inhibitors

If β-lactam sensitivity in *Mtb* is truly a consequence of classical drug resistance (i.e. truly collateral), we expect that genes/proteins implicated in β-lactam sensitivity should have close biochemical and/or regulatory connections to those that are implicated in classical drug resistance. We hypothesised that such connectivity may be detected as differential expression of β-lactam^s^ genes in response to classical drug treatment. Therefore, we investigated the differential expression of 111 genes with reported involvement in β-lactam sensitivity in *Mtb* or the closely related species *M. segmantis* or *M. bovis* (β-lactam^s^ genes, see Supp. Table S1) in response to incubation of *Mtb* with classical TB drugs (EMB, ETH, two FLQ (levofloxacin and ofloxacin), AMI, SM, INH, PZA and RIF) data [16]. Since we aimed to find commonalities between drug treatments, gene expression across single agent drug treatments were pooled.

We found that β-lactam^s^ genes showed a greater variability of expression than non-β-lactam^s^ genes (Kolmogorov-Smirnov (KS) test p-value 0.0034, Wilcoxon test p-value 0.00047, see Fig. 1A), indicating that classical drug treatment disproportionately affects the activity of these genes. We further validated this result by assessing the variability of randomly selected subsets of non-β-lactam^s^ genes matching the number of β-lactam^s^ genes (10,000 permutations, p=0.0023).

**Fig 1.**
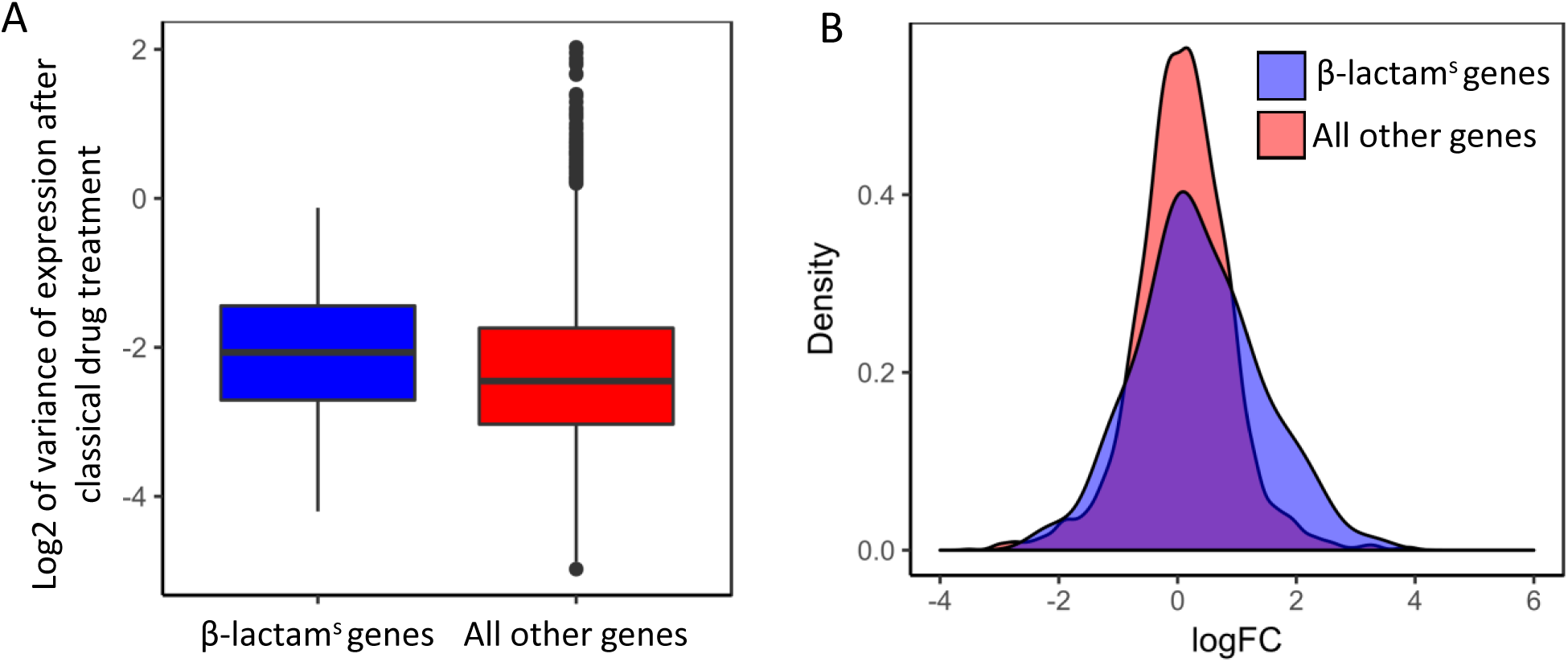
The expression of β-lactam^s^ genes is affected by treatment with classical TB drugs. **A.** β-lactam^s^ genes tend to be more variable than the rest of genes, suggesting these genes are affected by drug treatment (KS test p-value 0.00338). **B.** Log fold change (logFC) of β-lactam^s^ genes and all other non- β-lactam^s^ genes after classical drug treatment. β-lactam^s^ genes tend to have a more positive logFC than other genes, suggesting preferential activation.

We next performed differential expression analysis using limma [17], revealing 804 differentially expressed genes after drug treatment (q-value < 0.05, |fold change (FC)| > 2) (Supp. Table S3). We found that 31.46% of these were β-lactam^s^ genes (28/89, p = 0.0010 by Fisher exact test), of which 21 (75.0%) were upregulated (Fig. 1B). To check that the activation of β-lactam^s^ genes was not simply due to a general stress response, we also calculated the differential expression after incubation in an acidic stress environment (pH 4.8 to 5.6), which has been shown to restrict the growth of *Mtb* [28]. We found that only 5.41% of differentially expressed genes under acidic stress were β-lactam^s^ genes (29/536), indicating that drug treatment may be more likely to lead to differences in expression of β-lactam^s^ genes than other conditions that restrict cell growth.

Interestingly, *blal* (Rv1846c), the major repressor of the *blaC* β-lactamase, as well as *atpH* (Rv1307) and *sigC* (Rv2069), members of the BlaI regulon (Sala et al., 2009), were all upregulated after classical drug treatment (logFC = 1.509, q-value = 0.0095; logFC = 1.173, q-value = 0.03; logFC= 1.432, q-value = 0.0004, respectively). These data indicate that classical TB drug treatment may inhibit the main β-lactamase responsible for *Mtb’s* intrinsic β-lactam resistance. Among the upregulated genes we also found Rv1884c (*rpfC*) (logFC = 1.734, q-value = 0.0324), which has also been associated with β-lactam sensitivity [29]. Overall, treatment of *Mtb* with classical anti-TB drugs used in the clinic promoted the upregulation of key inhibitors of β-lactam resistance.

### 2. Strong co-regulation between β-lactam^S^ and DR genes

The findings from our gene expression analyses suggested that β-lactam^s^ genes may be co-regulated with those encoding the classical drug targets. Therefore, we searched for co-expression associations between the β-lactam^s^ and DR genes, the latter of which includes those that encode the classical drug target proteins.

First, we investigated module co-membership of β-lactam^s^ genes with DR genes among previously defined coexpression modules derived from 437 perturbation experiments with different drugs and growth-inhibitory conditions ([16], Supp. Table S2). We found that 47.37% of co-expression clusters containing DR genes also included at least one β-lactam^S^ gene, suggesting that these genes are controlled by similar regulatory networks.

Next, we compared the strength of correlation of expression of DR genes with β-lactam^s^ genes in these perturbation experiments (see Methods). We found that the majority of DR genes (27/37, 72.97%) had a stronger correlation with the genes of the β-lactam^s^ cluster than with all other genes (genes located above the diagonal line, see Fig. 2). These DR genes were disproportionately associated with INH resistance (10 genes) or to resistance to multiple drugs (7 genes), suggesting that these DR genes likely exert a strong regulatory influence on genes associated with β-lactam sensitivity.

**Fig 2.**
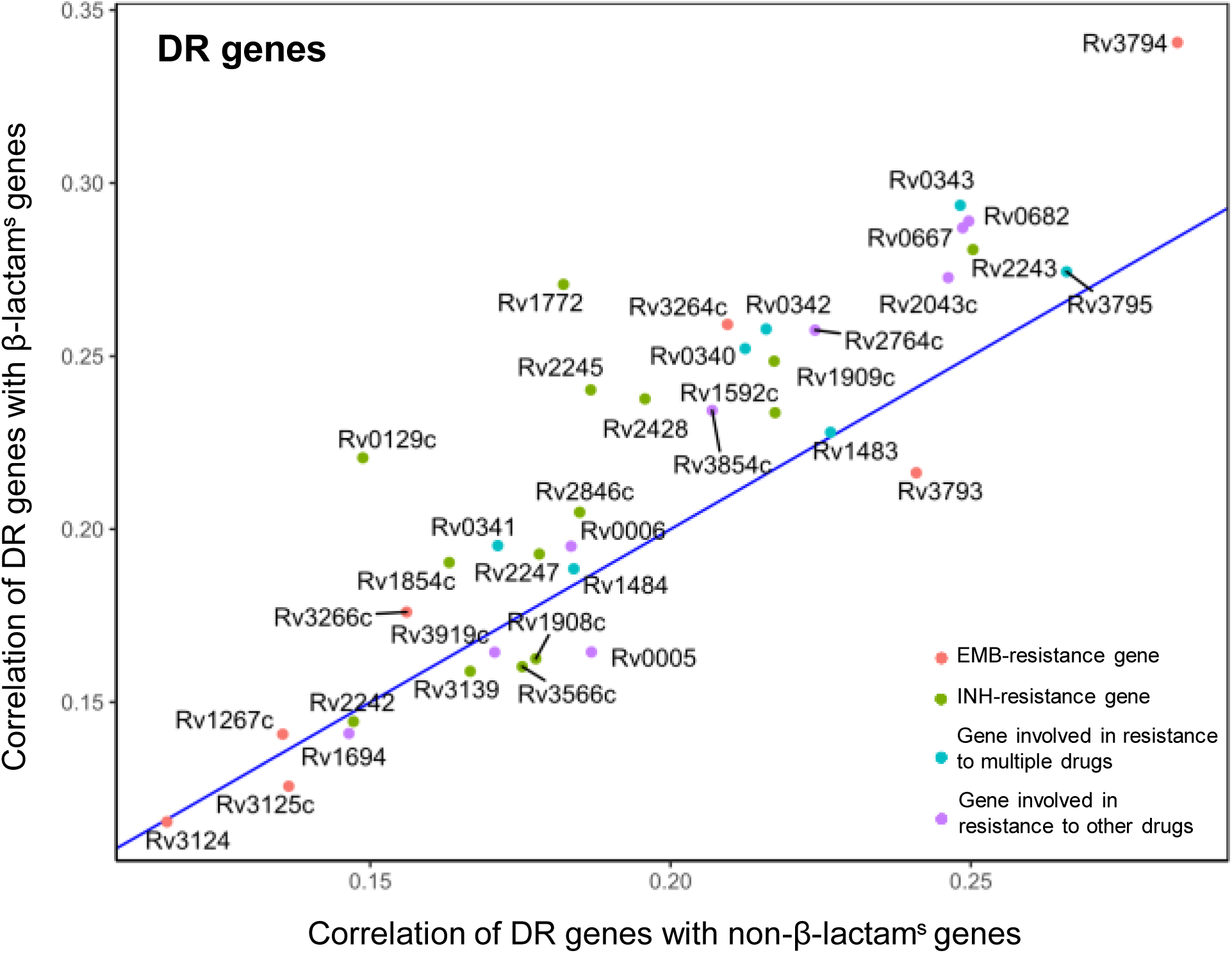
Upperquantile of expression correlation of DR genes with β-lactam^s^ genes (y-axis) and non-β-lactam^s^ genes (x-axis). Genes over the diagonal line are more strongly co-expressed with β-lactam^s^ genes than with other genes, and vice versa. The genes with the strongest positive correlation of expression (well above the diagonal line), especially *Rv2846c*, *Rv2243*, *Rv2245*, *Rv2247*, *Rv0129c*, are implicated in INH resistance.

Overall, DR and β-lactam^s^ genes were found to be highly co-expressed in the transcriptional network of TB, supporting our hypothesis of a co-regulatory association of these genes in *Mtb*.

### 3. β-lactam^S^ and DR nodes (genes/proteins) are highly linked in the molecular network of *Mtb*

To determine whether the co-expression associations between β-lactam^s^ and DR genes were the result of direct co-regulation between these genes as opposed to indirect associations, we investigated their localization and interaction in the *Mtb* protein-protein and gene regulatory networks. We integrated the STRING database [18] and transcription factor-target gene data published in [19], excluding duplicated edges and self-loops. The resulting network contained 4181 nodes (genes/proteins) and 37,313 edges, including experimentally validated physical and transcription factor – target associations between genes or proteins. There was no evidence to suggest that the combined PPI and GR network was not drawn from a power-law distribution (*p* = 0.065, i.e. indicating that we cannot reject the null hypothesis that the degree distribution follows a power-law distribution), supporting the view that the network structure is consistent with a true biological network [30]).

We found that the majority of β-lactam^S^ nodes (47 of 57) were localized in a highly specific network region (shown in Fig. 3). Network analysis using link and node permutation in BinoX revealed that β-lactam^S^ nodes are more likely to interact with each other than expected by chance (q-value < 1×10^−30^ using link and node permutation of the PPI network and q-value = 2.82×10^−5^ and q-value = 1 using link and node permutation, respectively). Interestingly within this subnetwork, β-lactam^s^ nodes were clustered based on the gene/protein functional role (Fig. 3), with well-defined clusters of nodes representing genes/proteins involved in metabolism, cell cycle and peptidoglycan biosynthesis, consistent with previous findings that gene function is related with network localization [31, 32]. However, the broader clustering of β-lactam^s^ nodes suggests a high degree of association between these genes even when these are functionally highly varied (Supp. Table S1), suggesting involvement in similar protein complexes or enzymatic reactions.

**Fig 3.**
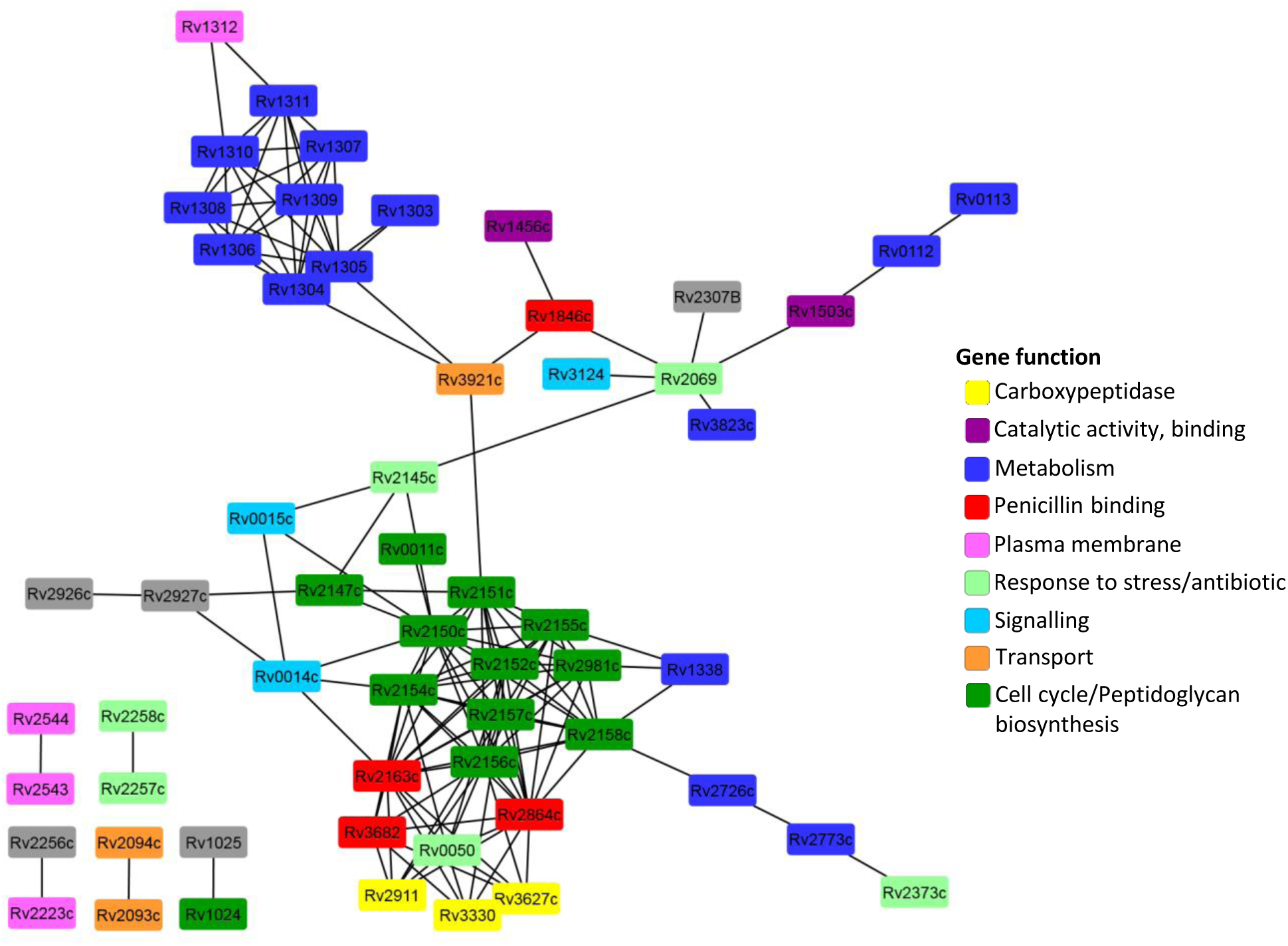
Network of interactions between genes/proteins associated with β-lactam sensitivity. β-lactam^S^ genes form a single interconnected network, with a few exceptions (lower left), indicating a high degree of localization in the global *Mtb* network. Nodes are coloured by predicted functional categories. The network shown is a combination of the PPI and the GR networks.

Next, we investigated the interactions between β-lactam^S^ nodes and DR nodes (Fig. 4, Supp. Table 4). We noted that the β-lactam^S^ nodes were located near the core or centre of the subnetwork, with DR nodes organized in clusters at the periphery grouped by drug type. Using BinoX, we found a significant crosstalk in the GR network between EMB resistance and β-lactam^S^ genes (q-value = 0.014 for link permutation and q-value = 0.065 for node permutation), and in the PPI network between INH and β-lactam^S^ genes (q-value = 0.009 for link permutation and q-value = 0.01 for node permutation). These results, together with the strong co-expression association between β-lactam^S^ and DR genes, supports our hypothesis of a direct co-regulation between β-lactam^s^ genes/proteins and the genes/proteins implicated in resistance to at least two of the first-line treatments used to treat TB in the clinic.

**Fig 4.**
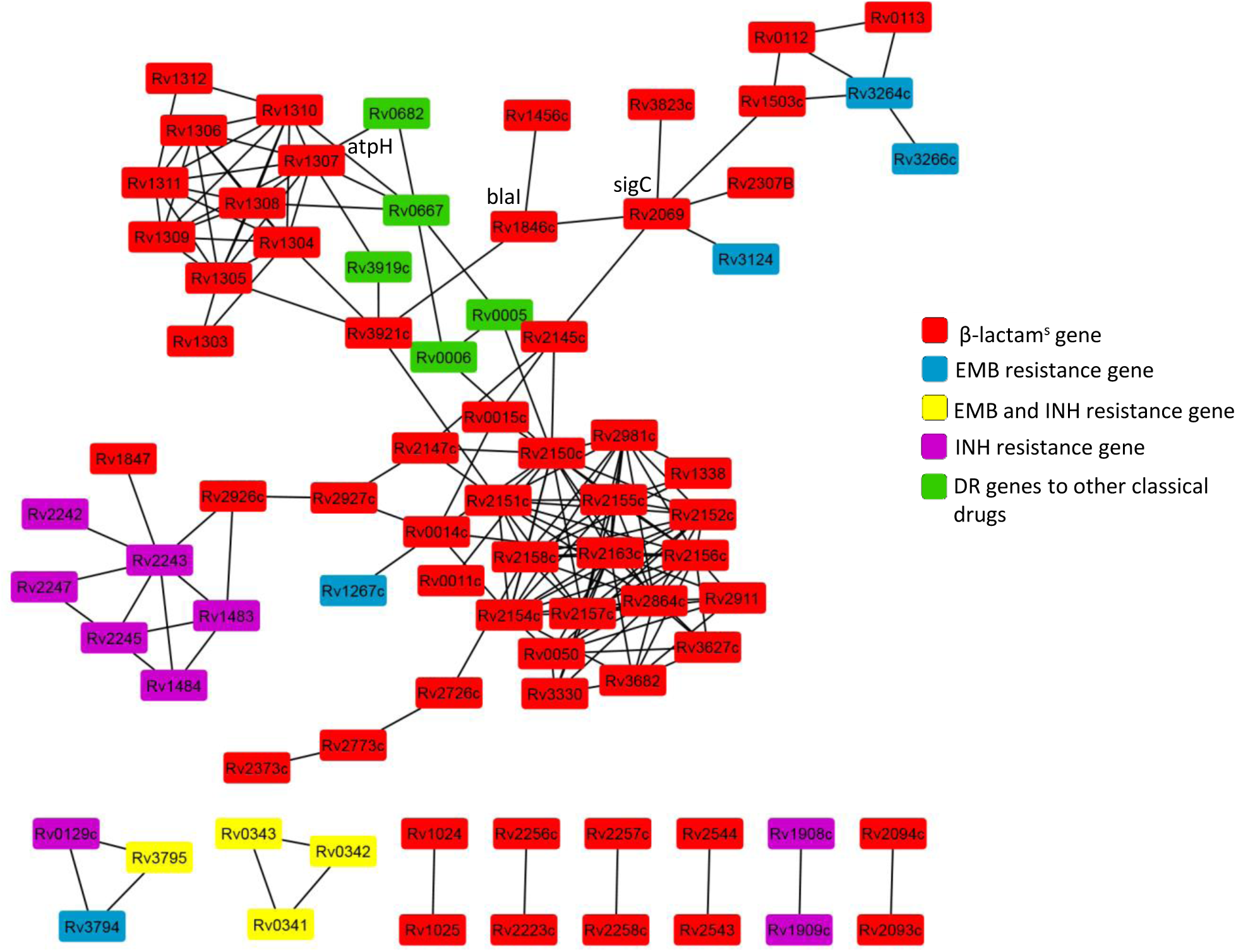
Network of interactions between β-lactam^S^ and DR genes/proteins. β-lactam^S^ genes/proteins tend to be located towards the core of the network, connecting distinct subgroups of DR genes. The network shown is a combination of the PPI and the GR networks.

### 4. *atpH* and *sigC* are key mediators of the interactions between DR and β-lactam^S^ genes

To identify the key genes linking β-lactam^S^ and DR genes/proteins, which likely mediate collateral β-lactam sensitivity, we performed random walks between the β-lactam^S^ and DR nodes in the PPI and GR networks. We calculated the access times between DR and β-lactam^S^ nodes in the GR and PPI networks, which represent the degree of influence between pairs of nodes (ie. a measure of ease of information flow between nodes, where lower scores suggest that biological information such as signalling or regulation is more likely to transfer from one gene/protein to another).

Since the importance of a node highly depends on the underlying network structure, we first determined how the structural differences between the PPI and GR networks would affect the random walks. We considered both directions of information flow, from DR nodes to β-lactam^S^ nodes and vice versa. We ranked the access times between all pairs of nodes in each database separately, and used bivariate Spearman’s ρ to calculate the similarity of stability scores [24, 25] (Supp. Fig. S1). We found that only the top 6.76% (272/4,019) of access time ranks in the DR to β-lactam^S^ node direction and 29.87% (1,146/3,836) in the β-lactam^S^ to DR node direction were consistent between networks. This result is consistent with the notion that PPI networks and GR networks represent different types of associations between genes and proteins. Therefore, we used PPI and GR networks separately for subsequent random walk analyses.

Ranking pairs of β-lactam^s^ and DR nodes by their access times revealed discrete sets of node pairs with similar influence (Fig. 5A-D), consistent with the modular organization of the PPI and GR networks [33]. We selected the set of node pairs with the smallest access times (red lines define threshold for each case); 30 and 35 pairs derived from the PPI and GR networks respectively in the DR to β-lactam^S^ direction; 160 and 103 pairs, respectively in the opposite direction (Supp. Table S5). These sets represent pairs of β-lactam^s^ and DR genes/proteins that are likely to modulate or influence each other’s activity.

**Fig 5.**
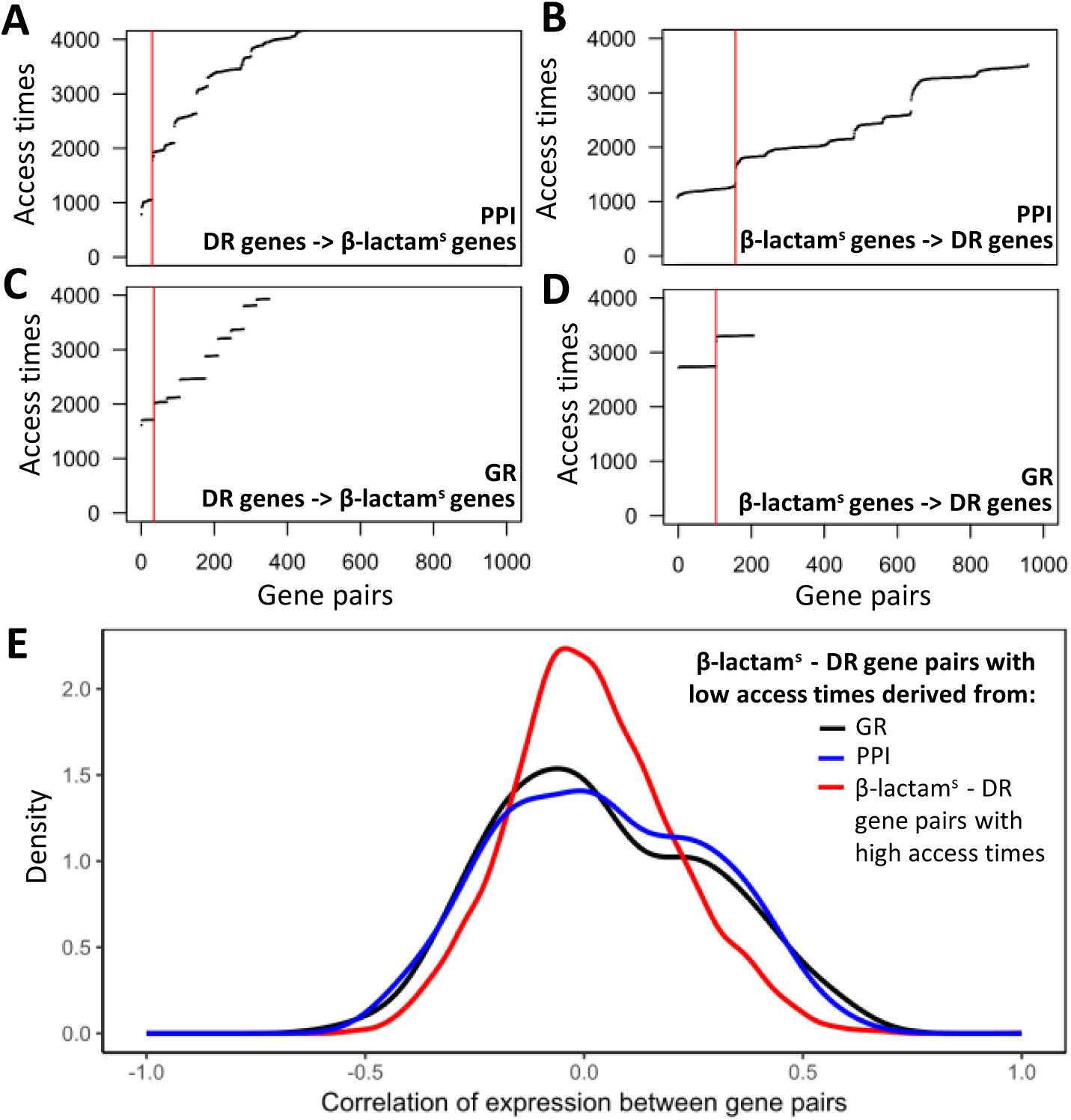
Highly influential pairs of β-lactam^s^ and DR nodes identified by random walks in the networks. **A-D.** Access times for gene pairs in the PPI (**A**, **B**) and GR (**C**, **D**) networks in the DR-β-lacam^s^ direction (**A**, **C**) and β-lactam^s^ -> DR direction (**B**,**D**). Due to a modular network structure, discrete clusters of highly influential pairs are identified. The set of genes with smallest access times (high influence between each other) to the left of the vertical red lines were used in subsequent analysis. (**E**) Strength of co-expression between DR-β-lactam^s^ node pairs. DR-β-lactam^s^ node pairs with lowest access times are more strongly correlated than pairs of genes with higher access times (wider distribution) in both the GR (KS test p=0.01) and PPI (KS test p = 0.0024) networks.

To ensure that the set of gene pairs with smallest access times was biologically meaningful, we compared the coexpression of pairs of DR-β-lactam^S^ genes with the smallest access times to pairs of DR-β-lactam^S^ genes with higher access times (Fig. 5E). Of note, the distributions were significantly different (Kolmogorov-Smirnov test p = 0.0024 in the PPI network, p-0.0024 in the GR network), with the distribution of co-expression of DR-β-lactam^S^ genes with the smallest access times being wider than the reference distribution, indicating an enrichment of stronger magnitudes of co-expression between these pairs.

By examination of DR-β-lactam^S^ gene pairs with the smallest access times we identified two key nodes in the paths of information flow: All low access time pairs derived from the PPI network were centred around AtpH (encoded by *atpH*, *Rv1307*), and those derived from the GR network were centred around *sigC* (*Rv2069*). Both *atpH* and *sigC* are transcriptionally regulated by BlaI [34], which is an important inhibitor of the *blaC* β-lactamase gene. This result once more implicates *blaI* and its transcriptional network in *Mtb* β-lactam collateral sensitivity.

### 5. *In silico* functional validation of a dependence mechanism between β-lactams^s^ and DR gene pairs

Finally, we sought to validate the functional association between β-lactams^s^ and DR genes/proteins by exploring their role in cell growth. We simulated the growth effects of β-lactams^s^ and DR gene pair knockouts using an *in silico* regulatory model, that incorporates both transcriptional data and metabolic modelling [26]. We found that simultaneous knockout of DR and β-lactams^s^ gene pairs caused a marked reduction of growth rate (growth rate < 0.010) or resulted in cell death more often than expected by chance (83.1% of DR-β-lactams^s^ pairs vs. 39% other pairs, Fisher exact test *p* = 3.25e-05, Fig. 6A, B), suggesting synthetic lethality and functional dependency between these genes. Of note, we found synthetic lethality between each of the *sigC*, *atpH* and *blaI* genes with the key DR genes *embB*, *katG* and *furA* (Fig. 6C).

**Fig. 6.**
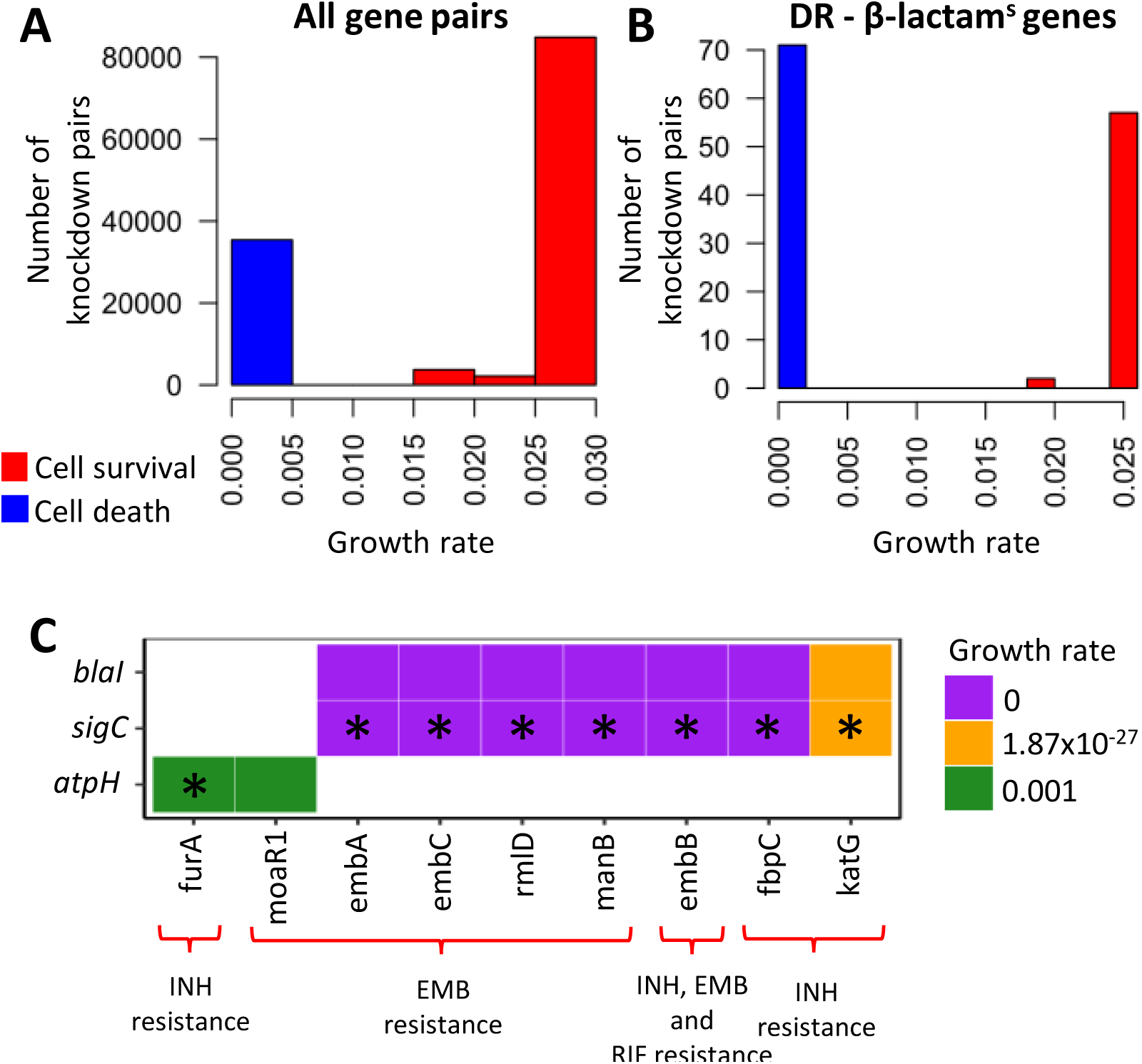
*In silico* double knockout of DR and β-lactam^s^ gene pairs reduce *Mtb* growth rate. Effect on the growth of *Mtb* after *in silico* knockout of all gene pairs (**A**) or DR-β-lactam^s^ gene pairs (**B**). DR-β-lactam^s^ gene pairs are enriched in those that lead to lethality (growth rate < 0.01) after knockout (Fisher test *p* =3.1 ×10^−9^). Knockouts resulting in cells with a growth rate < 0.010 were considered as lethal (blue), and above this cutoff, non-lethal (red). **C.** Growth rate of pairs of β-lactam^s^ and DR genes after double knockouts. The knockout of *blaI*, *atpH* and *sigC* in combination with DR-genes implicated in the resistance to commonly used drugs (e.g. EMB and INH) leads to cell death. Gene pairs identified by random walk network analyses as being highly influential pairs are indicated with an asterisk.

## Discussion

Here we demonstrate a novel systems biology approach to the investigation of *Mtb* β-lactam collateral sensitivity. We combined gene expression and network analyses and show that the inhibitor of intrinsic β-lactam resistance, *blaI*, is activated after treatment with classical anti-TB drugs (e.g. isoniazid, rifampicin, amikacin, streptomycin, levofloxacin, ofloxacin, ethambutol, ethionamide, pyrazinamide). Two genes transcriptionally regulated by *blaI*, *atpH* and *sigC* [34], as well as *Rv1884c* (*rpfC*), whose knockout mutants suffer increased sensitivity to β-lactams [29], were also upregulated. These findings support a model whereby classical anti-TB treatment drives cells towards a loss of β-lactam resistance, consistent with previous reports that drug-resistant *Mtb* were more likely to be susceptible to β-lactam treatment [7, 8].

Our co-expression analysis indicated a tight co-regulatory association between DR and β-lactam^s^ genes. Subsequent analysis of *Mtb* molecular networks supported these findings and identified a number of direct linkages. Together with evidence of strong gene co-expression associations, this finding suggests direct coregulation as opposed to indirect associations (as discussed in [35]).

Previous works have demonstrated the utility of random walks across networks to identify putative treatment cotargets for *Mtb* [13]. Here we applied random walks to identify key mediators of the communication between DR and β-lactam^s^ genes, and identified *atpH* and *sigC* as key regulators. In addition, *in silico* growth models revealed synthetic lethality after simultaneous knockout of any of *blaI*, *atpH* and *sigC* in combination with the genes conferring resistance to isoniazid, ethambutol or rifampicin, further supporting a functional association between these gene classes.

Our results point towards a model of collateral β-lactam sensitivity in classical drug resistant *Mtb*, involving a concerted effect of multiple genes. Others have also recently found that collateral sensitivity to β-lactams, mainly penicillins, develops in *Mtb* strains evolved *in vitro* to be resistant to the classical drug class aminoglycosides [9]. Our results suggest that *blaI*, together with its downstream targets, *atpH* and *sigC*, is a key regulator of collateral sensitivity resulting from classical drug resistance, although we were not able to detect a direct effect on transcription of the *blaC* β -lactamase gene in these data. Nevertheless, our evidence supporting a strong transcriptional wiring between β-lactam^s^ genes and DR genes suggests a tight co-evolutionary relationship, likely due in part to functional similarities between the genes, many of which are implicated in resistance to drugs that target *Mtb* cell wall biosynthesis e.g. ethambutol and isoniazid [36]. Thus, collateral sensitivity to β-lactams may represent a functional evolutionary trade-off to classical drug resistance.

The development of bacterial drug resistance is often accompanied by a fitness cost [37], which in some cases can be overcome by compensatory mechanisms. We speculate that β-lactam sensitivity arises in *Mtb* as a compensatory mechanism to regain fitness after disruption of the molecular network of TB due to the evolution of classical drug resistance. Indeed, genes associated with sensitivity to β-lactams (e.g. *murE*, *ponA1*, *murD*, *Rv2752c*, *Rv1218c*) have been identified as being under convergent evolution in resistant *Mtb* or harbouring compensatory mutations [38-40]. Although most studies have associated compensatory mechanisms with mutations [38-40], our results suggest that transcriptional changes might also be playing a role i.e upregulation of *blaI*. This assertion is consistent with a recent report showing that gene expression changes was associated with an increased fitness in *Mtb* that had developed resistance to rifampicin, isoniazid, streptomycin, fluoroquinolone, ethionamide and amikacin during a single patient infection [41].

Taken together, our findings implicate a potential role for β-lactam therapy in patients with classical drug-resistant TB, or the cyclic use of β-lactams with classical treatments to delay and/or prevent the development of resistance. Previous *in vitro* studies have demonstrated anti-TB activity for certain β-lactams plus β-lactamase inhibitor combinations [6] and other drugs [42]. However, mixed success in the clinic [11, 12, 43] suggests that treatment effectiveness might be dependent on other factors, such as the genetic background of the strain. Consequently, it will be essential to continue to develop our understanding of this phenomenon such that we can readily identify strains and therefore patients for whom β-lactam therapy may be appropriate, e.g. through processes akin to precision medicine in cancer treatment [44]. Our results suggest that subsets of patients with drug-resistant tuberculosis are more likely to benefit from β-lactam treatment.

## Conclusions

By integrating network analysis and gene expression data in a novel, systems-biology context we show that collateral β-lactam sensitivity in *Mtb* is driven through transcriptional regulation mediated through a small number of key interactions. In addition, our findings lend further support for exploration of combination anti-TB treatments that include β-lactams, particularly for patients infected with classical drug resistant TB.

## List of Abbreviations

TB: tuberculosis
*Mtb*: *Mycobacterium tuberculosis*
DR: drug resistance
β-lactam^s^: β-lactam sensitivity genes
GR: Gene regulatory
PPI: Protein-protein interaction
RIF: rifampicin
INH: isoniazid
AMI: aminoglycosides
SM: streptomycin
FLQ: fluoroquinolones
EMB: ethambutol
ETH: ethionamide
PAS: para-aminosalisylic acid
PZA: pyrazinamide
KS: Kolmogorov-Smirnov
FC: fold change

## Acknowledgements

We thank David Goode and Danielle Ingle for critical reading of the manuscript and Natalie Gunn for her support throughout this project.

